# Novel therapies for cancer-induced bone pain

**DOI:** 10.1101/2024.02.19.580951

**Authors:** Rayan Haroun, Samuel J. Gossage, Federico Iseppon, Alexander Fudge, Sara Caxaria, Manuel Arcangeletti, Charlotte Leese, Bazbek Davletov, James J. Cox, Shafaq Sikandar, Fraser Welsh, Iain P. Chessell, John N. Wood

**Author notes:** **Correspondence:** John N. Wood.

## Abstract

Cancer pain is a growing problem, especially with the substantial increase in cancer survival. Reports indicate that bone metastasis, whose primary symptom is bone pain, occurs in 65-75% of patients with advanced breast or prostate cancer. We optimized a preclinical *in vivo* model of cancer-induced bone pain (CIBP) involving the injection of Lewis Lung Carcinoma cells into the intramedullary space of the femur of C57BL/6 mice or transgenic mice on a C57BL/6 background. Mice gradually reduce the use of the affected limb, leading to altered weight bearing. Symptoms of secondary cutaneous heat sensitivity also manifest themselves. Following optimization, three potential analgesic treatments were assessed; 1) single ion channel targets (targeting the voltage-gated sodium channels Na_V_1.7, Na_V_1.8, or acid-sensing ion channels), 2) silencing µ-opioid receptor-expressing neurons by modified botulinum compounds, and 3) targeting two inflammatory mediators simultaneously (nerve growth factor (NGF) and tumor necrosis factor (TNF)). Unlike global Na_V_1.8 knockout mice which do not show any reduction in CIBP-related behavior, embryonic conditional Na_V_1.7 knockout mice in sensory neurons exhibit a mild reduction in CIBP-linked behavior. Modified botulinum compounds also failed to cause a detectable analgesic effect. In contrast, inhibition of NGF and/or TNF resulted in a significant reduction in CIBP-driven weight-bearing alterations and prevented the development of secondary cutaneous heat hyperalgesia. Our results support the inhibition of these inflammatory mediators; and more strongly their dual inhibition to treat CIBP, given the superiority of combination therapies in extending the time needed to reach limb use score zero in our CIBP model.

## 2. Introduction

Cancer-induced bone pain (CIBP) is the most common symptom in patients with bone metastases (1). Reports indicate that bone metastasis occurs in 65-75% of patients with advanced breast or prostate cancer and approximately 30-40% of patients with advanced lung cancer (2). Bone-metastasizing cancers cause pain due to the interplay between peripheral and central-related factors. Amongst the peripheral factors, acidosis is believed to be a major component, as bone metastasizing cancers result in excessive osteoclast activation, which subsequently leads to acidosis. Acidosis is known to cause pain, as sensory neurons contain a plethora of acid-sensing ion channels and TRPV1 transient receptor potential channels. Additionally, tumor cells and their associated stromal cells release mediators that are linked to pain, and these include nerve growth factor (NGF) and tumor necrosis factor alpha (TNFα). Moreover, research has highlighted the role of neuropathic pain in CIBP, leading to peripheral and central sensitization (3). We, therefore, investigated three potential modalities to treat CIBP: 1) single ion channel targets, namely voltage-gated sodium channels (Na_V_1.7 and Na_V_1.8) and acid-sensing ion channels 2) silencing µ-opioid receptor-expressing neurons using modified botulinum compounds and 3) dual targeting of tumor-derived products (NGF and TNFα).

The first approach was to target voltage-gated sodium channels and acid-sensing ion channels. Several pathophysiological studies have highlighted the role of enhanced expression and/or enhanced conductance of voltage-gated sodium channels as the final pain-causing mechanism for many mediators released in CIBP. For example, Qiu et al. (4) showed that the expression of Na_V_1.8 and Na_V_1.9 increases significantly in Sprague-Dawley rats bearing 256 mammary gland carcinoma cells in the intramedullary space of the tibia. Similarly, the activity of Na_V_1.8 channels can be modulated by several inflammatory mediators indirectly via their effects on protein kinases that phosphorylate Na_V_1.8 (5–7). NGF, which is considered a hallmark of CIBP, was shown to increase the expression of Na_V_1.7 and Na_V_1.8 and can allow the opening of these channels to take place at more negative membrane potentials (8–10). These studies show that voltage-gated sodium channels could play a crucial role in CIBP, and therefore, a significant part of this work focused on these channels. The first voltage-gated sodium channel targeted here is Na_V_1.7. This channel has gained huge interest because loss of function mutations in the *SCN9A* gene, which encodes for Na_V_1.7 channels, lead to pain insensitivity in humans (11). The role of this channel in pain is also supported by gain of function mutations in the *SCN9A* gene which are linked to erythromelalgia (12) and pain conditions like paroxysmal extreme pain disorder (13). Several animal models have been generated to study the role of Na_V_1.7 in pain sensation. The global Na_V_1.7 knockout mice die soon after birth, presumably due to the inability to feed (14). Therefore, conditional knockout models were generated, and results from these mouse models have shown that Na_V_1.7 is a critical player in acute pain sensation and chronic pain models (15, 16). The second channel studied here is Na_V_1.8, which has been shown to play a role in inflammatory and neuropathic pain conditions (17–20). Looking at CIBP specifically, it was shown that ablating or silencing Na_V_1.8+ neurons reduces pain-like behavior associated with CIBP in mice (21), and blocking (22) or downregulating (23) this channel resulted in an analgesic effect in rat models of CIBP. Therefore, we were also interested in checking whether congenital Na_V_1.8 knockout mice have reduced pain-like behavior after CIBP. We also looked at acid-sensing ion channels, mainly because mediators released in CIBP can lead to increased expression of acid-sensing ion channels (ASICs) (24) and because local acidosis is a proposed factor that contributes to CIBP (3). To target ASICs, we relied on mambalgin-1. Mambalgin-1 is a 57-amino acid peptide found in African black mamba venom that blocks subtypes of acid-sensing ion channels. Analgesia obtained after mambalgin application is comparable to that obtained following morphine administration (25). What makes mambalgins an interesting therapeutic option is that they lack toxicity, and their analgesic effect does not depend on the opioid system (25).

The second approach utilized here is to silence µ-opioid-expressing neurons by modified botulinum compounds as a longer-lasting and potentially safer alternative to opioid therapy. Small doses of morphine are administered intrathecally as part of a successful treatment option for refractory cancer pain. While efficacious, it was also reported that this treatment option affected the day-to-day activities of cancer patients and is linked to tolerance (26). Additionally, previous work indicated that chronic morphine treatment exaggerates CIBP, bone loss, and spontaneous fracture in a mouse model of bone cancer (27). Therefore, alternative means of targeting the opioid system could be beneficial. Botulinum neurotoxin serotype A is composed of a light-chain zinc endopeptidase as well as a heavy chain that binds to neuronal receptors and promotes light-chain transfer across the endosomal membrane (28). Once incorporated within the cell, the light chain can inhibit neurons for many months by cleaving synaptosomal-associated protein 25 (SNAP25), an important protein for neurotransmitter release (29–31). Maiarù et al. (32) leveraged a recently established “protein stapling” technique (29, 31) that enabled the nonchemical linking of recombinantly generated proteins, employing core components of the SNARE (soluble N-ethylmaleimide–sensitive factor attachment protein receptor) complex, to achieve irreversible linking of two distinct peptide segments into a functional unit (30). This group used SNARE proteins to bind the light-chain domain of botulinum neurotoxin type A (BOT) to dermorphin, which targets µ-opioid receptor-expressing neurons. This construct renders the BOT specific for the µ-opioid receptor-expressing neurons, solving the protein toxicity problems related to botulinum-based compounds. After binding to the µ-opioid receptors, the construct is internalized due to the presence of the translocation domain. Subsequently, the protease domain of the toxin is released into the cytoplasm, causing a profound suppression of neurotransmitter release. This group found that the intrathecal injection of 100 ng of this construct to mice resulted in significant analgesia following neuropathic pain and inflammatory pain models (32). The authors postulated that the analgesic target for this construct did not depend on the µ-opioid receptor-positive primary afferents but rather the µ-opioid receptor-positive dorsal horn neurons (32). To assess its therapeutic potential in CIBP, Derm-BOT was used to silence the µ-opioid receptor-expressing neurons.

The third approach was to target two tumor-derived inflammatory mediators simultaneously. The two targets chosen were NGF and TNFα. NGF can be produced by various types of cancer cells in the bone (e.g., those derived from primary tumors of the breast and prostate). On the other hand, TNFα is one of the cytokines contributing to pain in bone cancer models as not only cancer cells, but also immune cells (including macrophages, leukocytes, and thrombocytes), tend to release it in CIBP (33). NGF can lead to the sensitization of sensory neurons directly by increasing the expression of ion channels linked to nociceptive signal transduction and transmission (such as TRPV1, ASICs, and voltage-gated sodium channels) through its effects after binding TrkA receptors. NGF also triggers the release of inflammatory mediators (including histamine, serotonin, and more NGF) from immune cells (34, 35). Due to the transcriptional changes caused by NGF-mediated TrkA activation, neurotransmitters like substance P, calcitonin gene-related peptide (CGRP), and brain-derived neurotrophic factor are also increased. The binding of all these peptides to their receptors on the second-order neuron together with glutamate may result in intense depolarization of the postsynaptic receptors of the second-order neurons, increasing the probability of N-methyl-D-aspartate (NMDA) receptor activation, enhancing the probability of central sensitization. Therefore, NGF is implicated not only in peripheral inflammation but also in increasing the excitability of primary afferents by causing various transcriptional changes (24). The usefulness of targeting NGF in CIBP is supported by clinical (36) and preclinical findings (37–39). Regarding TNFα, it contributes to heat and mechanical hyperalgesia in CIBP (40–45). Inhibiting the effects of TNFα could be achieved by using mAbs against it or its soluble receptor, TNFR1 (46, 47). In this report, MEDI578 was used to inhibit NGF, and etanercept, which can also bind TNFβ, was used to inhibit TNFα.

## 3. Materials and Methods

### 3.1 Cell culture

Lewis lung carcinoma (LL/2) cells (from American Type Culture Collection (ATCC)) were cultured in a medium containing 90% Dulbecco’s Modified Eagle Medium (DMEM) and 10% fetal bovine serum (FBS) and 0.1% Penicillin/Streptomycin for 14 days before the surgery. DMEM was supplemented with L-glutamine (1%) and glucose (4.5 g/liter). The cells were sub-cultured whenever ∼80% confluency was reached, which was done a day before the surgery. On the surgery day, LL/2 cells were harvested by scraping and were centrifuged at a speed of 1500 rpm for two mins. The supernatant was removed, and the cells were resuspended in a culture medium that contained DMEM to attain various final concentrations: ∼2×10^7^ cells/ml, 2×10^6^ cells/ml, or 2×10^5^ cells/ml for the CIBP model optimization experiments and ∼2×10^6^ cells/ml (resulting in the inoculation of ∼2×10^4^) for all the remaining studies. The cell counting and viability check were carried out using the Countess Automated Cell Counter (Thermo Fisher Scientific).

### 3.2 Mice

Mice were housed in groups of 2–5 per cage with a 12-hour light/dark cycle and were allowed free access to water and a standard diet. Mice were acclimatized for two weeks before the surgery, and the baseline measurements were taken at the end of these two weeks. All experiments were performed with the approval of personal and project licenses from the United Kingdom Home Office according to guidelines set by the Animals (Scientific Procedures) Act 1986 Amendment Regulations 2012 and guidelines of the Committee for Research and Ethical Issues of IASP.

### 3.3 Breeding strategies for transgenic mice

#### 3.3.1 Conditional Na_V_1.7 knockout mice

To generate conditional Na_V_1.7 knockout mice in the DRG neurons, heterozygous mice expressing Cre under the control of the advillin promoter (identifier B6.129P2-Avil^tm2(cre)Fawa^/J, (48)) and homozygous floxed for Na_V_1.7 (identifier Scn9a^tm1.1Jnw^, (14)) were crossed. Primers used to genotype Na_V_1.7 flox mice and Advillin-Cre mice can be found in (49).

#### 3.3.2 Na_V_1.8 knockout mice

Homozygous global Na_V_1.8 knockout mice were crossed with heterozygous Na_V_1.8 knockout mice (identifier Scn10a^tm1Jnw^, (17)). Resulting homozygous Na_V_1.8 knockout mice were to test the effect of channel knockout by comparing them to the heterozygous littermates (control). Heterozygous null mutant Na_V_1.8 mice were used as controls since they were shown to be phenotypically identical to wild-type mice (14). Genotyping for Na_V_1.8 knockout mice can be found in (50).

### 3.4 Surgery

Surgery was carried out on anesthetized mice (males and females). Anesthesia was achieved using 1.5-3% isoflurane. The legs and the thighs of the mice were shaved, and the shaved area was cleaned using hibiscrub solution. A sterile lacri-lube was applied to the eyes. The reflexes of the mice to pinches were checked to ensure successful anesthesia. An incision was made in the skin above and lateral to the patella on the left leg. The patella and the lateral retinaculum tendons were loosened to move the patella to the side and expose the distal femoral epiphysis. A 30G needle was used to drill a hole through the femur to permit access to the intramedullary space of the femur. The 30G needle was removed, and a 0.3ml insulin syringe was used to inoculate ∼2×10^4^ LL/2 cells suspended in DMEM (for the optimization studies, a range of LL/2 cell numbers was tested). The hole in the distal femur was sealed using bone wax (Johnson & Johnson). To ensure that there was no bleeding, the wound was washed with sterile saline. Following that, the patella was re-placed into its original location, and the skin was sutured using 6–0 absorbable vicryl rapid (Ethicon). Lidocaine was applied at the surgery site, and the animals were placed in the recovery chamber and monitored until they recovered. The choice of the animal number was based on previous experience from animal experiments in our lab.

### 3.5 Compound administration and doses

For the NGF/TNF studies, MEDI578 (MedImmune, 3mg/kg) and/or etanercept (Pfizer, 10mg/kg) were administered intraperitoneally on day 10 after surgery. The control group for this study included 13 mice that were treated with PBS and 10 mice treated with NIP-228 (isotype control antibody for MEDI578) at a 3mg/kg dose. All compounds were dissolved so that each mouse received 10µl of the diluted biologic per gram of its body weight. For mambalgin-1 testing, mambalgin-1 (Smartox) was dissolved in PBS and was used at a dose of 34µM via intrathecal injection. When morphine was administered intrathecally, a 60ng dose was used, and when administered subcutaneously a 15mg/kg dose was used. Morphine used in this study was used in the form of morphine sulfate (Hamlen pharmaceuticals). For the Derm-BOT study, mice received an intrathecal injection of 100ng of Derm-BOT (or the molar equivalent of the control (unconjugated BOT)) (32). Derm-BOT and unconjugated BOT were a kind present from Professor Bazbek Davletov (University of Sheffield, UK).

### 3.6 Behavioral tests

After the surgery, the limb-use score of mice was checked daily. Any score less than four on the sixth day after the surgery caused the mouse to be sacrificed and excluded from the study. This rule ensured that the observed pain phenotype was only due to the cancer growth and not a complication from the surgery.

#### 3.6.1 Limb-use score

The mice housed in the same cage were placed together in a glass box (30 × 45 cm) for at least five minutes. Then each mouse was left in the glass box on its own and was observed for ∼4 minutes, and the use of the ipsilateral limb was estimated using the standard limb use scoring system in which: 4 indicates a normal use of the affected limb; 3 denotes slight limping (slight preferential use of the contralateral limb when rearing); 2 indicates clear limping; 1 clear limping, and with a tendency of not using the affected limb; and 0 means there is no use of the affected limb. Reaching a limb-use score of zero was used as an endpoint for the experiment.

#### 3.6.2 Static weight-bearing

The weight-bearing test was done as previously described by (21) and was reported as a percentage of the total weight on both paws.

#### 3.6.3 Hargreaves’ test

The Hargreaves’ test was used to measure secondary cutaneous heat hyperalgesia and was performed as previously described by (51).

#### 3.6.4 Analysis of the median time needed to reach limb-use score zero

The time needed for a mouse to reach zero limb-use score was considered an endpoint, and any mouse that reached this score was sacrificed. Therefore, the time needed to reach this score is considered an indirect pain measure, when the number of cancer cells injected is similar between mice. It is important to note that mice were also sacrificed if the tumor caused the mice to lose 15% of their body weight measured from the day of the surgery.

### 3.7 Micro-computed tomography (µCT)

After each mouse was sacrificed (at limb use score zero), its femurs, both ipsilateral and contralateral, were dissected. Femurs were then fixed using a 4% paraformaldehyde (PFA) solution for 24 hours. Then, PFA was removed, and femurs were washed using phosphate-buffered saline (PBS). Femurs were then kept in 70% ethanol (v/v) at 4°C until scanned using μCT.

### 3.8 Statistical analyses

The mixed-effects model (REML) was used with multiple comparisons to compare the pain-like behavior between two groups over time. In all statistical tests, the difference between groups is considered significant when the p-value is <0.05. All statistical analyses were performed using GraphPad Prism 10. Data are presented as mean ± standard error of the mean (SEM). For comparing the time needed to reach limb-use score zero, the Log-rank (Mantel-Cox) and the Gehan-Breslow-Wilcoxon tests were used, and the median duration to reach this limb-use score was compared between groups. The Dunnett’s test for multiple comparisons was used to compare the mean results after the surgery with the baseline readings. To compare three (or more) groups with each other, the one-way analysis of variance (ANOVA) test was used, followed by multiple comparisons. Statistical significance is expressed as: *p < .05; **p < .01; ***p < .001, ****p < .0001.

## 4. Results

### 4.1 CIBP model optimization

Previous published papers from our lab studying CIBP in C57BL/6 mice or transgenic mice on a C57BL/6 background relied on injecting 2×10^5^ LL/2 cells in the intramedullary space of the femur. In this model, the median time needed to reach limb-use score zero was short (less than 20 days). Therefore, we attempted to reduce the number of LL/2 cells injected to increase the time needed to reach limb-use score zero and slow down the pain phenotype progression in order to make a model that is more representative of chronic pain conditions. Therefore, two lesser cancer cell numbers were attempted, 2×10^4^ (red, Figure 1) and 2×10^3^ (blue, Figure 1), and they were compared to the original cancer cell number (black, Figure 1). The reduction of the number of cancer cells injected significantly increased the study duration and showed a dose-dependent reduction in pain phenotype intensity. By referring to the limb scoring results (Figure 1 A), it appears that the mean limb score becomes significantly lower than the baseline readings after 10 days for the 2×10^5^ cells group, 15 days for the 2×10^4^ cells group, and 19 days for the 2×10^3^ cells group. Weight-bearing reduction followed similar dose-dependent trends (Figure 1B).

**Figure 1:**
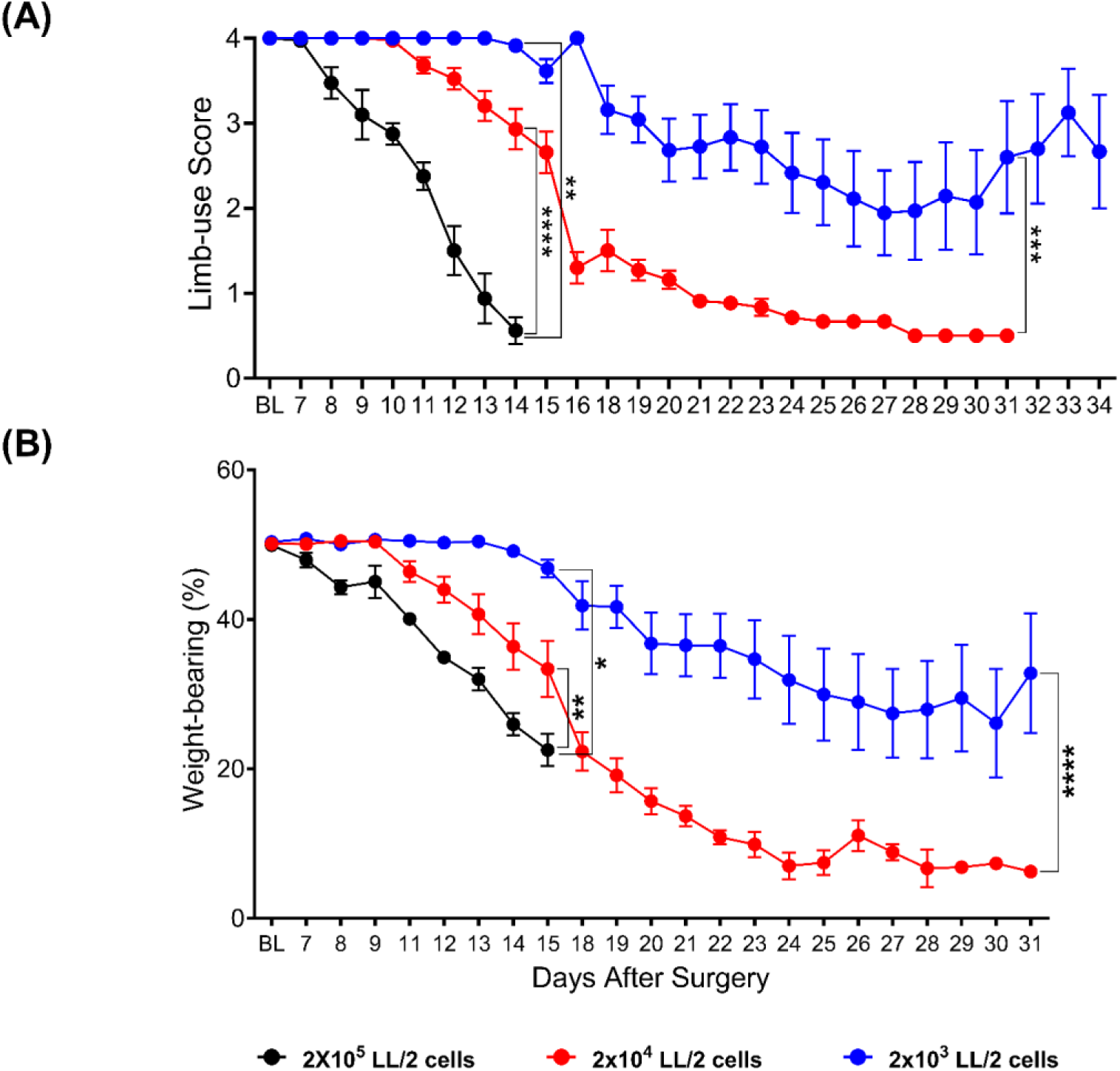
The injection of LL/2 cells into the intramedullary space of the femur of mice caused a significant decline in the limb-use score (A) and the weight-borne on the ipsilateral limb (B). This decline was dependent on the number of LL/2 cells injected. (A) The mean limb-use score became significantly lower than the baseline on day 10 for the 2×10^5^ cancer cells group (p-value= 0.0095), day 15 for the 2×10^4^ cancer cells group (p-value 0.0007), and day 19 for the 2×10^3^ cancer cells group (p-value 0.0349). (B) The analysis of the weight-bearing results showed that the reduction in the weight-bearing became significant (compared to the baseline) on day 11 after surgery for the 2×10^5^ group (p-value= 0.0444), on day 12 after the surgery (p-value= 0.0456) in the mice that received 2×10^4^ cells and after 14 days in the mice that received 2×10^3^ cancer cells (p-value 0.0212). Analyses of the limb-use scoring, and weight-bearing results showed that the effect of the cancer cells is statistically significant, with the speed of reduction of limb-use and weight-borne on the ipsilateral limb being directly proportional to the number of cancer cells inoculated (asterisks of significance are shown on the graph (the mixed-effects model)). At the baselines, n=12 for the 2×10^4^ group and the 2×10^3^ group (equal mix of sexes), and n=5 males for the 2×10^5^ cancer cells group. REML analysis (followed by Dunnett’s test for multiple comparisons) was used to estimate the time required to cause a significant drop in either the weight-bearing or the limb-use score compared to the baseline after inoculating a specific number of cancer cells in the femur.

Because decreasing the number of cancer cells injected slowed the speed of the pain phenotype progression, we decided to reduce the number of cancer cells in our subsequent experiments. Between the two lesser cancer cell numbers (2×10^4^ and 2×10^3^ cells), 2×10^4^ cells was selected to be used for the following experiments as it showed less variability between mice. Additionally, in the group injected with 2×10^3^ LL/2 cells two (out of 12) mice did not develop any pain-like behavior until the end of the study, i.e., day 40 after surgery. µCT analyses indicated that injecting 2×10^4^ LL/2 cells into the intramedullary space of the femur of C57BL/6 mice caused a clear bone degradation (Figure 2).

**Figure 2:**
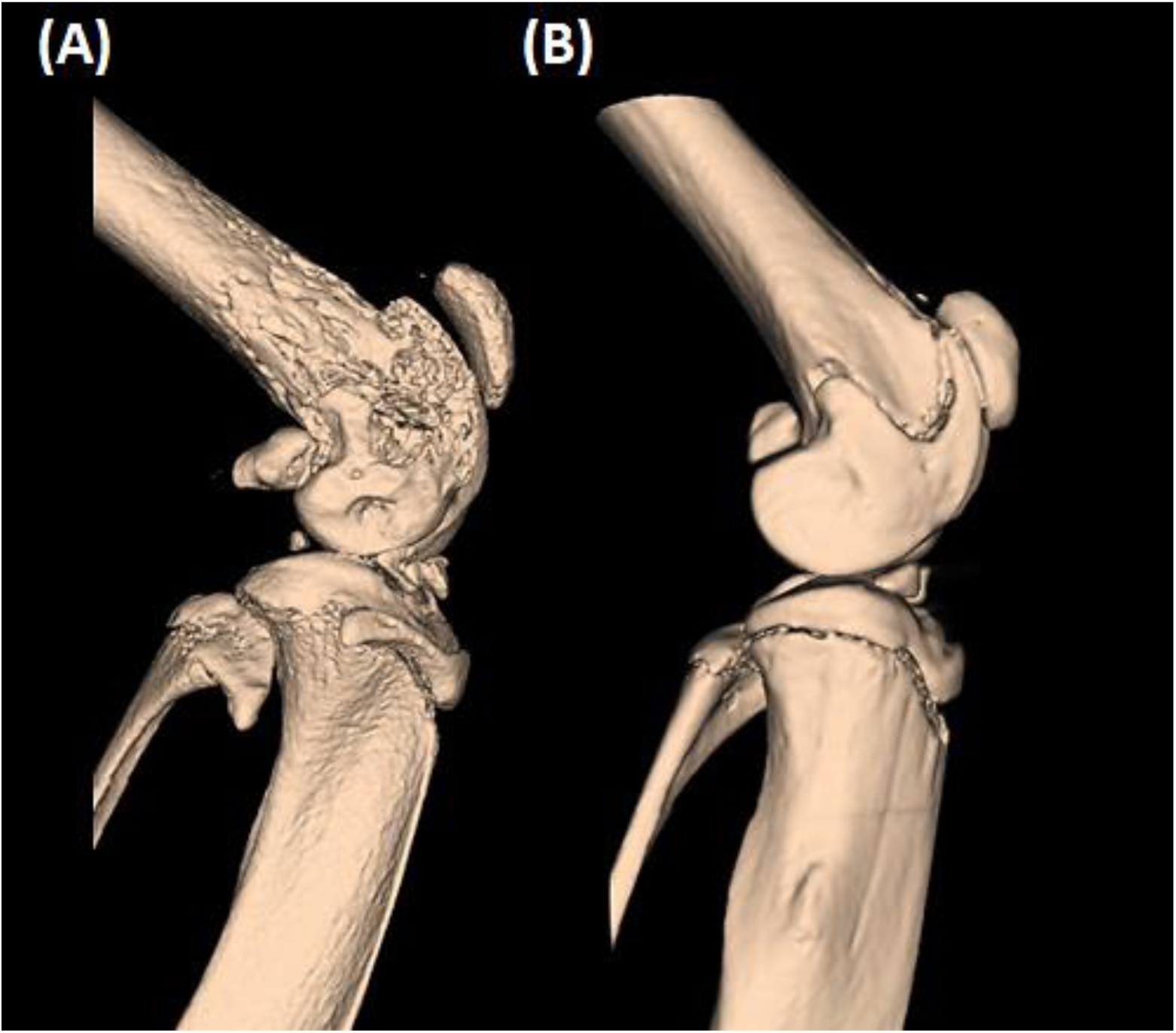
µCT imaging showing osteolysis induced by the growth of LL/2 cells in the femurs of C57BL/6 mice. The tumor growth causes bone degradation in the ipsilateral femur (A), leaving the contralateral femur (B) unaffected.

### 4.2 Single ion channel targets in CIBP

#### 4.2.1 Voltage-gated sodium channels and CIBP

##### 4.2.1.1 Na_V_1.7 conditional knockout mice show a modest reduction in pain-like behavior associated with CIBP

Previous research demonstrated that several mediators derived from cancer cells and their associated stromal cells sensitize and/or increase the expression of Na_V_1.7 channels in rodents (8–10, 52). Therefore, the role of this channel in CIBP was tested by assessing whether mice in which this channel is conditionally knocked out in DRG neurons could show a reduction in the pain phenotype caused by bone cancer. Therefore, conditional Na_V_1.7 knockout mice (and Cre-negative littermates) underwent CIBP surgery to inoculate LL/2 cells into their left femurs. The Na_V_1.7 conditional knockout mice exhibited modestly better mean limb-use scores compared to their littermates (Figure 3A (i)) and showed a slower reduction in the weight-borne on the ipsilateral limb (Figure 3A (ii)).

**Figure 3:**
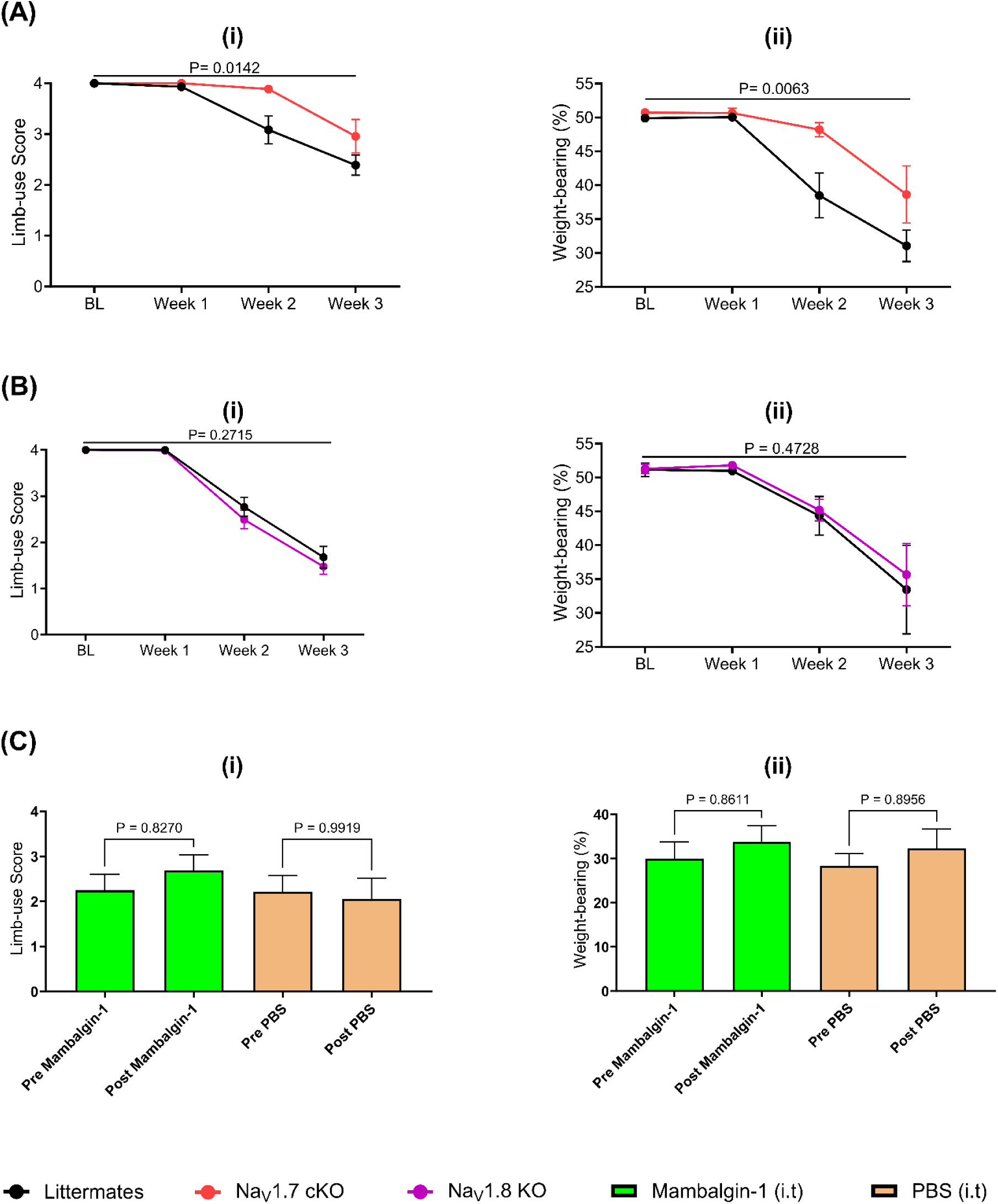
An investigation of the potential significance of Na_V_1.7 channels, Na_V_1.8 channels, and acid-sensing ion channels as targets to treat CIBP. A) Conditional Na_V_1.7 knockout mice in the DRGs showed a modest reduction in pain phenotype associated with CIBP. (A (i)) A comparison of the limb-use scores between the conditional Na_V_1.7 knockout mice (Na_V_1.7 cKO) and their littermates in the CIBP model. Results indicate that the decline in the use of the affected limb in the Na_V_1.7 cKO mice was significantly slower than that of the control group (the REML, p-value=0.0142). (A (ii)) Ipsilateral weight-bearing after the CIBP surgery in Na_V_1.7 cKO mice and their littermates. The REML analysis revealed that the difference in the weight-bearing between the two groups was statistically significant (p-value=0.0063), with the Na_V_1.7 conditional knockout mice showing a significantly less reduction in the percentage of weight put on the ipsilateral limb over time compared to their littermates. N=9 in the Na_V_1.7 cKO group (five females and four males) and 14 for their littermates (seven males and seven females). B) Knocking out the expression of Na_V_1.8 channels congenitally did not cause a statistically significant alteration in the pain phenotype associated with CIBP when assessed using limb-use score (B (i)) and weight-bearing (B (ii)). N=8 for littermates (two males and six females) and 12 for Na_V_1.8 knockout mice (Na_V_1.8 KO, six males and six females). C) The intrathecal administration of mambalgin-1 (34 µM) did not alter the limb-use score (C (i)) or weight-bearing of mice with CIBP significantly (C (ii)). Behavioral tests were performed 30 minutes after mambalgin-1 or phosphate-buffered saline (PBS) administration. N=10 for the mambalgin-1 group (five males and five females) and eight for the PBS group (four males and four females). For Figures A and B, data were analyzed using REML analysis, while for Figure C, data were analyzed using the One-way ANOVA test with multiple comparisons.

##### 4.2.1.2 Congenital global Na_V_1.8 knockout mice show a normal progression of pain-like behavior after CIBP

Previous work using neuronal ablation techniques and neuronal silencing tools indicated that Na_V_1.8+ neurons play a role in CIBP (21). Additionally, knocking down the expression of this channel reduced CIBP in rats (23). Accordingly, we decided to investigate whether deleting Na_V_1.8 channels congenitally could reduce CIBP in mice. Our results showed that Na_V_1.8 knockout mice developed similar pain-like behavior compared to their littermates in our CIBP model when tested using limb-use scoring (Figure 3B (i)) and weight-bearing (Figure 3B (ii)).

#### 4.2.2 Acid-sensing ion channels and CIBP

As previous work showed that mambalgins result in an analgesic effect comparable to morphine but with fewer side effects, we decided to use mambalgin-1 to block acid-sensing ion channels in our CIBP model. Our results indicate that the intrathecal administration of 34 μM of mambalgin-1 did not result in a significant alteration of pain-like behavior after CIBP compared to the vehicle (Figure 3C). Behavioral tests included limb-use scoring (Figure 3C (i)) and ipsilateral weight-bearing (Figure 3C (ii)).

### 4.3 Neuronal Silencing in CIBP

#### Intrathecal Derm-BOT did not cause a significant alteration in pain-like behavior after CIBP

For this study, mice underwent the CIBP surgery, and whenever a mouse reached limb-use score 3, it was injected intrathecally with a modified botulinum compound that specifically inhibits neurotransmission in the µ opioid receptor-expressing neurons (Derm-BOT). The control group for this study involved the use of a botulinum compound that cannot enter cells due to the absence of opioid targeting moeity, and this was also injected upon reaching limb-use score 3. The comparison between the two treatment groups highlighted no significant difference in the limb-use score (Figure 4A (i)) or weight-bearing (Figure 4A (ii)). To test whether opioids reduce pain-like behavior in this model, morphine (15 mg/kg) was administered subcutaneously to mice when they reached limb-use score 3, and results indicated that morphine improved their limb-use scores (Figure 4B (i)) and ipsilateral weight-bearing (Figure 4B (ii)), bringing them to normal levels. Similarly, we conducted a pilot study to assess whether the intrathecal injection of 60 ng of morphine could reduce pain-like behavior in this model and the results indicated that morphine improved the limb-use score (Figure 4C (i)) and weight-bearing of mice with CIBP (Figure 4C (ii)). It is noteworthy that morphine was administered when mice reached limb-use score 3, which is the same time point used for the Derm-BOT administration.

**Figure 4:**
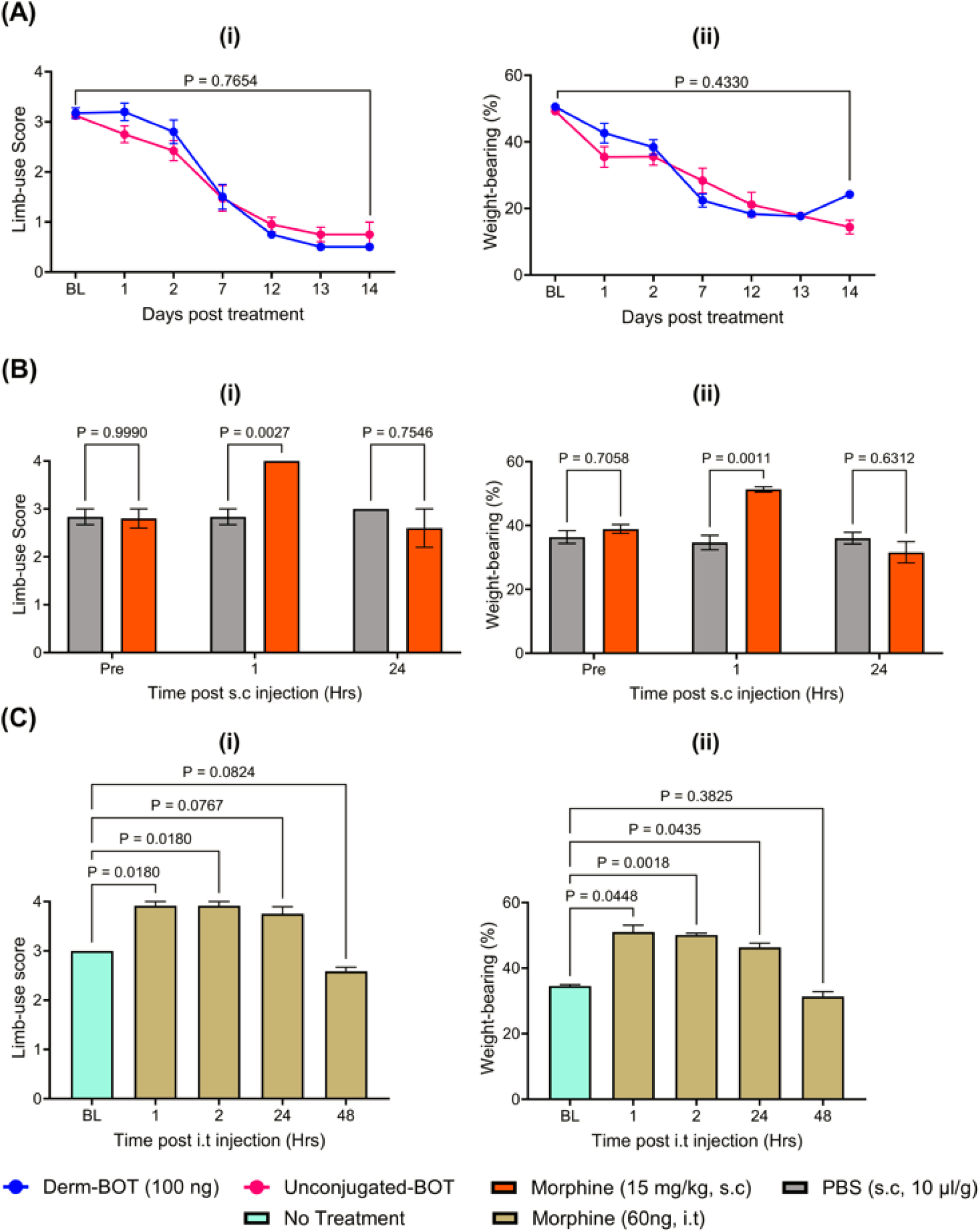
Attempts to silence µ-opioid receptor-expressing neurons in the spinal cord using modified botulinum compounds failed to reduce pain-like behavior in a mouse model of CIBP. Intrathecal and subcutaneous morphine administration resulted in significant analgesia. A) Modified botulinum compounds to silence the µ-opioid receptor-expressing neurons failed to reduce pain-like behavior in our mouse model of CIBP. Mice were assessed for recovery from surgery on day 7 after the surgery by measuring the limb-use score (shown as BL on graph (i)) and no mice showed signs of limping on that day. After that, the limb-use was scored daily using the standard limb-use scoring system. If a mouse reached limb-use score 3 (slight limping), it was injected intrathecally with either 100 ng of dermorphine botulinum (Derm-BOT, shown in blue) or the molar-equivalent amount of the control (un-conjugated BOT lacking dermorphine, shown in pink). Following the intrathecal treatment, several behavioral tests were performed, including the limb-use score (i) and the static weight-bearing (ii) tests. (A (i)) The comparison of the average limb-use score between the two treatment groups over time highlighted no significant difference between the two groups when assessed using the REML analyses (p-value for the treatment effect was 0.7654). (A (ii)) The comparison of the average percentage of the weight-borne on the affected limb between the two treatment groups over time indicated no significant difference between them when assessed using the REML analyses (p-value for the treatment effect was 0.4330). n=10 for each group at the baseline (five males and five females). B) Subcutaneous morphine (15 mg/kg) improved limb-use scores (i) and weight-bearing (ii) of mice with CIBP. Morphine was administered when mice reached limb-use score 3. Results were analyzed using repeated-measures one-way ANOVA test with multiple comparisons. N=5 (three males and two females) for the morphine group and six for the PBS group (three males and three females). C) Intrathecal morphine improved limb-use scores (i) and weight-bearing (ii) of mice with CIBP. Morphine was administered intrathecally at a dose of 60ng per mouse when mice reached limb-use score 3. Results were analyzed using repeated-measures one-way ANOVA test with multiple comparisons. N=3 (two males and one female). P-values for the one-way ANOVA tests are 0.0002 for the limb-use score test and 0.0083 for the weight-bearing test. The p-values for the multiple comparisons are shown in the figure.

### 4.4 Dual targeting of NGF and TNFα in CIBP

#### 4.4.1 The simultaneous inhibition of NGF and TNFα increased the median time needed to reach limb-use score zero after CIBP

As NGF and TNFα play essential roles in CIBP, we investigated whether blocking these two tumor-derived products could reduce the pain phenotype in our mouse model of CIBP (Figure 5). Etanercept (10 mg/kg) and MEDI578 (3mg/kg) were used to inhibit TNFα and NGF, respectively. The synergy between MEDI578 and Etanercept was evident as the combination was the only treatment option that resulted in a significant increase in median time needed to reach limb-use score zero after CIBP, compared to the control group (Figure 5A), with the median time needed to reach limb-use score zero in the control group being 12 days after treatment and 17 days in the combination-treated group. The median time needed to reach limb-use score zero in the etanercept-treated group was 13 days after the surgery, while that of the MEDI578-treated group was 15 days. Notably, these biologics were administered on day 10 after CIBP surgery via intraperitoneal injection. Behavioral differences are not thought to be driven by structural differences at the bone level because µCT analyses indicated that bone destruction was similar between treatment groups at the endpoint.

**Figure 5:**
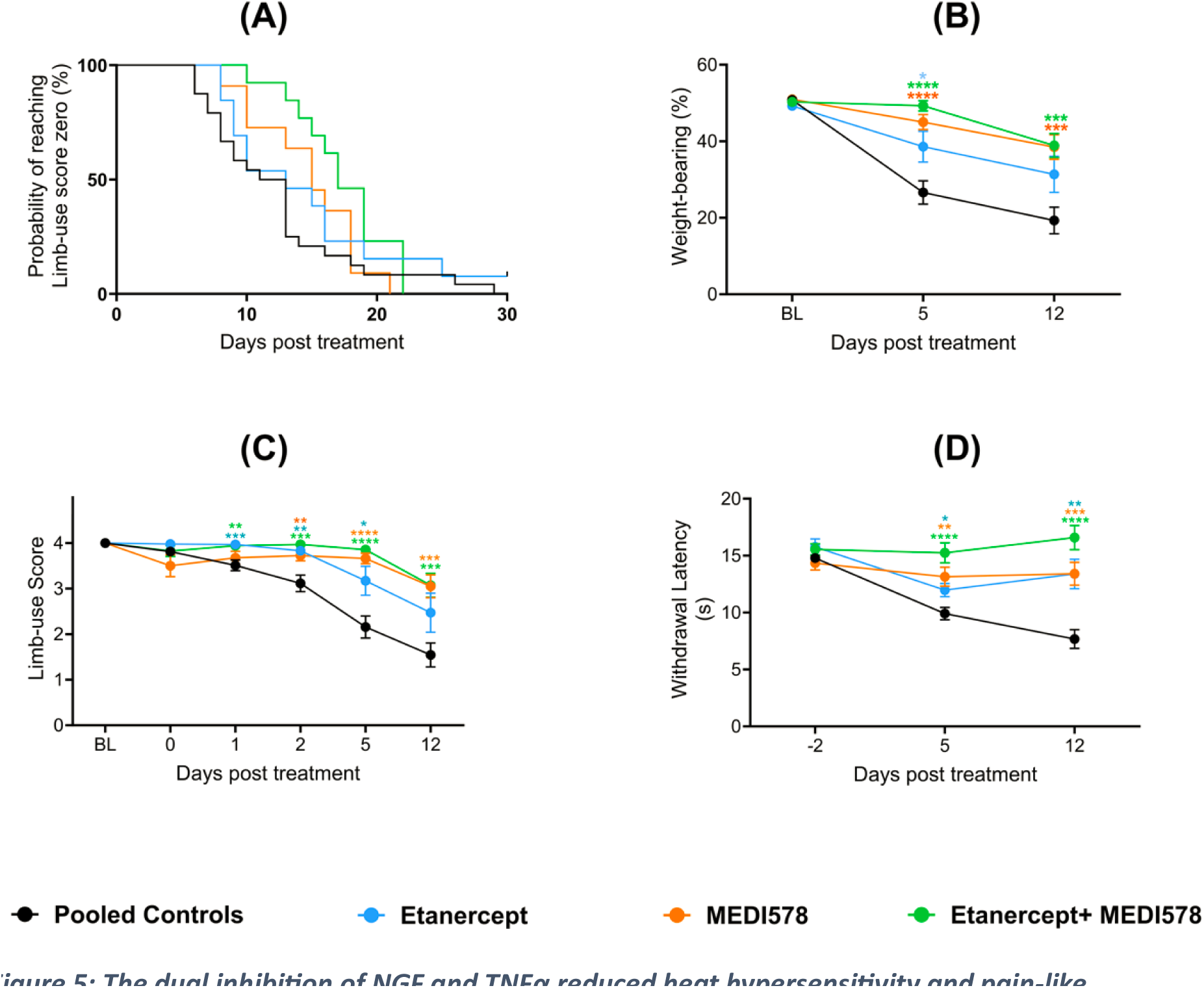
The dual inhibition of NGF and TNFα reduced heat hypersensitivity and pain-like behaviors associated with CIBP. A) Dual treatment using Etanercept and MEDI578 significantly increased the time needed to reach limb-use score zero of mice after CIBP surgery. The median time needed to reach limb-use score zero for the mice that received anti-NGF (MEDI578) + Etanercept (a protein that inhibits the binding of TNF to its receptors) was 17 days after treatment, while it was 12 days for the control group ((Gehan-Breslow-Wilcoxon test (p-value= 0.0026) and Log-rank Mantel-Cox test (p-value= 0.0357)). B) Antagonising the effects of NGF, TNF, or both slowed down the reduction in the weight put on the ipsilateral limb after CIBP. Compared to the control group, the etanercept-treated group had significantly higher mean weight-bearing results (the REML, 0.0113). Similarly, the mice treated with MEDI578 showed a significantly smaller drop in the weight-bearing (the REML, p-value<0.0001). The combination of MEDI578 and Etanercept also had a higher fraction of weight put on the ipsilateral limb compared to the controls (p-value=<0.0001). Mice treated with the combination of Etanercept and MEDI578 had a statistically significant higher weight-bearing performance than the ones treated with Etanercept alone (p-value=0.0098, the REML), but not compared to the mice treated with MEDI578 alone (p-value= 0.4159). C) Blocking the binding of NGF, TNF, or both to their receptors slowed down the reduction in the limb-use score associated with CIBP. Compared to the control group, the etanercept-treated group had a significantly higher mean limb-use score (the REML, p-value= 0.0024). Similarly, mice treated with MEDI578 showed better mean limb-use scores compared to control mice (the REML, p-value= 0.0029). The combination of MEDI578 and Etanercept also led to mice having better use of the affected limb compared to the controls (p-value<0.0001). Mice treated with Etanercept and MEDI578 did not significantly improve the limb-use score statistically compared to those treated with MEDI578 alone (p-value= 0.1673, the mixed-effects model) or Etanercept alone (p-value= 0.0693). D) The concurrent application of MEDI578 and Etanercept prevented the development of secondary cutaneous heat hyperalgesia associated with CIBP. The application of MEDI578 significantly diminished heat hyperalgesia associated with CIBP compared to the control group (p-value <0.0001). Similarly, Etanercept significantly lessened heat hyperalgesia associated with CIBP compared to the control group (p-value <0.0001). The simultaneous application of MEDI578 and Etanercept prevented the development of secondary cutaneous heat hyperalgesia entirely and had a significantly improved effect compared to Etanercept alone (p-value= 0.0097) and MEDI578 alone (p-value= 0.0137). The asterisks of significance shown in the figure represent the post hoc analysis results comparing each treatment with the control group at each time point. n=13 in each group at the baseline (except for the control group (n=23)). Doses: Etanercept (10mg/kg), MEDI578 (3mg/kg), and control antibody (3mg/kg). The drugs were injected intraperitoneally on day ten post-surgery. Control antibody (n=10, five males and five females), PBS (n=13, six males and seven females), MEDI578 (n=13, six females and seven males), etanercept (n=13, six males and seven females), and the combination (n=13, six females and seven males).

#### 4.4.2 Inhibiting NGF and/or TNFα slowed down the reduction of the use of the cancer-bearing limb and the weight-bearing behavior

The administration of Etanercept and/or MEDI578 reduced the pain-like behaviors associated with CIBP (weight-bearing (Figure 5B) and limb-use score (Figure 5C)) compared to the control group. The co-administration of MEDI578 and Etanercept always resulted in a more profound reduction in the pain phenotype compared to each treatment separately. This improvement was statistically significant compared to Etanercept alone in the weight-bearing test (p-value= 0.0098, REML, Figure 5B).

#### 4.4.3 Secondary cutaneous heat hypersensitivity is prevented by the co-administration of MEDI578 and Etanercept

The administration of Etanercept (a protein that inhibits TNF) and/or MEDI578 (anti-NGF) reduced secondary cutaneous heat sensitivity (Figure 5D) significantly compared to the control group when tested using the Hargreaves’ test. The combination treatment reduced the secondary cutaneous heat sensitivity to the extent that it reached statistical significance compared to Etanercept alone (p-value= 0.0097, REML). While the performance of MEDI578 was remarkable compared to the negative control and etanercept alone, it was still less effective than the combination treatment, as there had been a consistent trend of improved performance in the group treated with the combination. The superiority of the combination treatment compared to the use of MEDI578 alone was convincing in the Hargreaves’ test, with the REML analysis showing a p-value of 0.0137 when the two groups are compared.

## 5. Discussion

### 5.1 Single ion channel targets

The aim of this work was to identify targets for treating CIBP. Previous work from our lab and other labs reported potential treatment strategies for CIBP using mouse models (see Table 1). In this report, a mouse model of CIBP was optimized. This model involves the injection of ∼2×10^4^ Lewis Lung Carcinoma cells into the intramedullary space of the femur of C57BL/6 mice or transgenic mice on the C57BL/6 background. Static weight bearing and limb use score are both observed to decrease gradually as the model progresses (Figure 1). Previous work relied on injecting 2×10^5^ LL/2 cells in the femur, which allows mice to be studied for less than 20 days after surgery. When this model was employed by our lab to test whether knocking out Na_V_1.7 channels conditionally using Wnt-Cre could reduce pain associated with CIBP, results indicated a tendency for improved limb use in the Na_V_1.7 conditional knockout (Na_V_1.7^Wnt1^) mice compared to their littermates but without statistical significance (53). One potential reason for this could be that the CIBP model used was aggressive and did not allow a long enough pain window to assess the difference between groups. What further reinforces this hypothesis is our findings reported here, employing the refined model which involves injection of 2×10^4^ LL/2 cells (Figure 1). Here, we report modest but significant analgesia in the conditional Na_V_1.7 knockout group (Na_V_1.7^Adv^), indicating that Na_V_1.7 could be playing a role in CIBP (Figure 3A). With this model, we could observe mice for more than 30 days after surgery, which gave us a longer time to assess the analgesic potential of the therapeutic targets (Figure 1).

**Table 1:** Examples of studies reporting a significant reduction in pain-like behavior/hyperalgesia in mouse models of CIBP.

| **Analgesic target/strategy** | **Cancer cells** | **Behavioural tests** | **References** |
| --- | --- | --- | --- |
| **Ablation/Silencing Nav1.8+ neurons using chemogenetic tools** | LL/2 (lung cancer) | Static weight bearing | (21) |
| **Anti-programmed cell death-1 (anti-PD-1) monoclonal antibody** | LL/2, LL/2-Luc2, CMT 167 | Mechanical, thermal and cold sensitivity | (54) |
| **Pharmacologic inhibition of ferroptosis** | LL2-Luc | Mechanical and thermal sensitivity and spontaneous pain behaviour | (55) |
| **Melatonin evoked analgesia via sirtuin 1 (SIRT1)-dependent nucleus-high mobility group box-1 inhibition** | LL/2 | Mechanical and thermal sensitivity | (56) |
| **Inhibition of Wnt5b signalling using anti-Ryk antibodies** | NCTC2472 (osteosarcoma) | Mechanical and thermal sensitivity | (57) |
| **Blockade of spinal vascular endothelial growth factor A signalling using antibodies or receptor blockers** | LL/2 | Mechanical and thermal sensitivity | (58) |
| **miR-9-5p modified mouse bone marrow mesenchymal stem cell mediated analgesic actions** | NCTC | Spontaneous pain behaviour and mechanical sensitivity | (59) |
| **Srt 1720 activation of Sirtuin 1 relieves bone cancer pain** | NCTC2472 | Spontaneous pain behaviour and mechanical sensitivity | (60) |
| **Phosphatidylinositol 3-kinase (PI3K)-protein kinase B (Akt) inhibition** | Mouse 4T1 breast cancer cells | Mechanical sensitivity | (61) |
| **Opioids** | NCTC 2472 | Spontaneous lifting behaviour and Limb-use on rotarod | (62) |
| **Antibodies against nerve growth factor** | Canine prostate carcinoma | Ongoing nocifensive behaviors, mechanical | (63) |

|  |  | and thermal sensitivity |  |
| --- | --- | --- | --- |
| **Osteoprotegerin block of bone destruction** | NCTC2472 | Forced ambulatory guarding, limb-use score, spontaneous flinches and mechanical sensitivity | (64) |

Using the ‘refined cancer model’ we moved on to assessing more molecular targets. The second target tested was Na_V_1.8. Na_V_1.8 represents an appealing analgesic target, and several groups showed its role in inflammatory and neuropathic pain, and channel blockers were promising in phase II clinical trials in patients with painful diabetic peripheral neuropathy (65). Additionally, our previous work demonstrated that ablating/silencing Na_V_1.8+ neurons reduced CIBP. While data from channel knockdown studies on rats relying on the injection of Walker 256 breast tumor cells suggest that Na_V_1.8 could contribute to CIBP (23), knocking out Na_V_1.8 channels in mice did not lead to a significant change in CIBP compared to their littermates in our model (Figure 3B). The discrepancy in results could be due to species differences or compensatory mechanisms caused by congenital channel deletion as opposed to channel knockdown. To rule out the possibility of compensatory mechanisms, future work should look at knocking out the channel expression in adulthood.

The third target/s tested here focused on acid-sensing ion channels using mambalgin-1. While mambalgins block all acid-sensing ion channel combinations expressed in central neurons and nociceptors, we report these experiments as part of the single ion channel targets section as acid-sensing ion channels are not the sole acid sensors in the body. Therefore, blocking them is still subject to compensation by other targets, like TRPV1 channels, and in fact, the involvement of TRPV1 in sensing bone cancer-derived acidosis has been previously shown (3). What made mambalgins of great interest to us were *in vivo* mouse experiments that highlighted how significant analgesia is seen when mambalgins are administered (25). Because acidosis was suggested as a pain-causing mechanism in CIBP, we attempted to use mambalgin-1 as a potential analgesic in our mouse model of CIBP. Unfortunately, intrathecal mambalgin-1 (34µM) did not improve limb-use score or weight-bearing of mice with CIBP (Figure 3C). This could be attributable to the redundancy in pain pathways where blocking a single target can be easily compensated for by other targets. Because none of the single ion channel targets tested here seemed promising enough for therapeutic application, we decided to adopt broader analgesic approaches in our subsequent experiments, namely cell silencing and dual-targeting.

### 5.2 Cell silencing

While silencing µ-opioid receptor-expressing neurons using Derm-BOT could be a promising analgesic option that lessens problems associated with the intrathecal or systemic administration of opioid agonists (66), this approach failed to cause noticeable changes in the pain-like behavior in our mouse model of CIBP (Figure 4A). One limitation of our work is that it does not provide evidence for silencing µ-opioid receptor+ neurons by Derm-BOT, but our behavioral results clearly indicate that Derm-BOT did not cause any noticeable analgesic effects. Problems related to the target itself are unlikely because the subcutaneous (Figure 4B) and intrathecal (Figure 4C) administration of morphine caused analgesia. Because opioids resulted in a substantial analgesic effect in this work, one could attempt to use different doses of Derm-BOT to determine if another dose is needed to achieve a substantial lasting reduction in pain-like behavior. The dose used here was previously shown to reduce pain in various inflammatory and neuropathic models (32), but perhaps a different dose is needed to effect behavioral changes in CIBP.

### 5.3 Dual targeting of tumor-derived products

The hypothesis behind the inhibition of NGF is backed up by previous findings, which indicate that in CIBP, inhibiting NGF actions reduces neuromas (67), prevents central sensitization (68), and reduces the NGF-mediated neurotransmitters and ion channel expression elevation. Also, this target is validated by human genetics as loss of function mutations in the receptor TrkA are linked to pain insensitivity (69). More importantly, tanezumab (an anti-NGF antibody) resulted in a significant analgesic effect in clinical trials on patients with CIBP (36). Additionally, previous work from our lab found that ˃70% of bone afferents retrogradely labeled by injecting Fast Blue in the femur of C57BL/6 are Na_V_1.8+. Approximately 90% of Fast Blue+ Na_V_1.8+ neurons expressed TrkA receptors (70). On the other hand, inhibiting the binding of TNF to its receptor carries countless benefits in CIBP (71), like the prevention of secondary cutaneous heat and mechanical sensitivity.

The results indicate that this combination, unlike targeting a sole mediator (i.e. TNF or NGF), significantly increased the time needed to reach limb-use score zero after the CIBP surgery in C57BL/6 mice. Mice treated with etanercept and MEDI578 concurrently demonstrated significantly less pain-like behavior. Also, the combination-treated mice had significantly less secondary cutaneous heat hypersensitivity compared to those treated with MEDI578 or etanercept alone. It is worth highlighting the impressive phenotype achieved by using MEDI578 alone. This should not come as a surprise, given how previous work indicated the involvement of Na_V_1.8+ neurons in CIBP (21) and that more than 90% of bone afferents that are positive for Na_V_1.8 are also positive for TrkA (70). While these findings, coupled with the promising clinical trial results investigating the use of tanezumab in CIBP patients (36), suggest that targeting TrkA alone is an appealing method to treat CIBP, concerns related to the safety of using large doses of tanezumab remain compelling (72), especially because clinical trials indicated that doses needed for CIBP are significantly higher than those needed for osteoarthritis pain. The analgesic effects seen with MEDI578 could reignite the debate about the utility of single molecular targets for complex conditions. While it is evident that MEDI578 analgesic effects were remarkable, one could argue that when TrkA is activated/inhibited many downstream pathways/molecular targets are affected. Investigating the potential synergy between inhibiting NGF and TNF simultaneously would be beneficial not only from an efficacy point of view, but also, perhaps more importantly, from a safety viewpoint. To the best of our knowledge, this is the first study that assessed the analgesic efficacy of targeting NGF and TNF simultaneously in CIBP, but further work is needed to investigate the potential synergy. One option would be to test whether the coadministration of sub-efficacious doses of MEDI578 and etanercept could result in a significant analgesic effect in CIBP.

In conclusion, this work optimized a mouse model of CIBP by selecting an appropriate number of Lewis lung carcinoma cells for intra-femoral inoculation. Following that, various single ion channel targets were tested to investigate potential analgesic effects, namely Na_V_1.7 channels, Na_V_1.8 channels, and acid-sensing ion channels. While deleting Na_V_1.7 channels in sensory neurons showed a statistically significant reduction in pain-like behavior, the effect was only modest, especially when compared to the standard of care (i.e. opioids). These findings do not come as a surprise because drugs to treat pathologies of redundant biological systems often exhibit a high attrition rate in the clinic, mainly because the inhibition of one target can be easily compensated for by other targets.

While our model indicates that opioids, when administered intrathecally or subcutaneously, reduce pain-like behavior significantly, clinical data indicate their long-term use is associated with severe side effects, like tolerance, dependence, and respiratory depression. Preclinical CIBP models also suggested that sustained morphine accelerates sarcoma-induced bone pain, bone loss, and spontaneous fracture. Therefore, there is a pressing need to find alternative analgesic tools that are not only effective but also safe in the long term. Such novel analgesics are unlikely to be reliant on single ion channel targets given the system redundancy and the complexity of the condition.

Therefore, more generalized analgesic approaches are needed. Herein, we tested two approaches: silencing µ-opioid positive neurons in the spinal cord and using MEDI578 (to inhibit nerve growth factor) and Etanercept (to inhibit the tumor necrosis factor) simultaneously. While cell silencing did not show a good analgesic effect in our hands with the doses tested in this report, the simultaneous use of etanercept and MEDI578 resulted in an impressive reduction in the pain phenotype associated with CIBP and prevented the development of secondary cutaneous heat hyperalgesia. The analgesic potential of the combination of etanercept and MEDI578 was convincingly superior to the use of etanercept or MEDI578 alone. These findings, thus, open the door for the use of bispecific antibodies targeting these two mediators or low doses of each one of the two biologics to effect analgesia in patients living with CIBP.

## Author contributions

Conceptualization: RH, JJC, SS, JW

Methodology: RH, SG, FI, AF, SC, MA

Investigation: RH, SG, FI, AF, SC, MA

Funding acquisition: JW, SS

Project administration: SG, JW

Supervision: RH, FW, IC, JJC, SS, JW

Writing – original draft: RH, JW

## Acknowledgment

We would like to thank Professor Bazbek Davletov for providing Derm-BOT and its control. We would also like to thank Dr Iain Chessell and Dr Fraser Welsh for their kind donation of MEDI578 (and its control) and etanercept.

